# Genomic features underlie the co-option of SVA transposons as cis-regulatory elements in human pluripotent stem cells

**DOI:** 10.1101/2022.01.10.475682

**Authors:** Samantha M. Barnada, Andrew Isopi, Daniela Tejada-Martinez, Clément Goubert, Sruti Patoori, Luca Pagliaroli, Mason Tracewell, Marco Trizzino

## Abstract

Domestication of transposable elements (TEs) into functional cis-regulatory elements is a widespread phenomenon. However, the mechanisms behind why some TEs are co-opted as functional enhancers while others are not are underappreciated. SINE-VNTR-Alus (SVAs) are the youngest group of transposons in the human genome, where ∼3,700 copies are annotated, nearly half of which are human-specific. Many studies indicate that SVAs are among the most frequently co-opted TEs in human gene regulation, but the mechanisms underlying such processes have not yet been thoroughly investigated. Here, we leveraged CRISPR-interference (CRISPRi), computational and functional genomics to elucidate the genomic features that underlie SVA domestication into human stem-cell gene regulation. We found that ∼750 SVAs are co-opted as functional cis-regulatory elements in human induced pluripotent stem cells. These SVAs are significantly closer to genes and harbor more transcription factor binding sites than non-co-opted SVAs. We show that a long DNA motif composed of flanking YY1/2 and OCT4 binding sites is enriched in the co-opted SVAs and that these two transcription factors bind consecutively on the TE sequence. We used CRISPRi to epigenetically repress active SVAs in stem cell-like NCCIT cells. Epigenetic perturbation of active SVAs strongly attenuated YY1/OCT4 binding and influenced neighboring gene expression. Ultimately, SVA repression resulted in ∼3,000 differentially expressed genes, 131 of which were the nearest gene to an annotated SVA. In summary, we demonstrated that SVAs modulate human gene expression, and uncovered that location and sequence composition contribute to SVA domestication into gene regulatory networks.

## Introduction

Transposable elements (TEs) are mobile DNA sequences that account for over 50% of the human genome, yet there is very limited knowledge on the extent of their impact on genome evolution, function, and disease.

Many elegant studies have proposed that TE sequences constantly reshape eukaryotic gene regulation [1–24] yet the underlying mechanisms are largely uncharacterized. Several TE types are active and replication competent in humans, and the genomic dispersal of these elements can affect the regulatory configurations of proximal host genes. For example, TE insertions may introduce novel cis-regulatory elements (CREs = enhancers, promoters, insulators) at the gene locus [8,13,14,16,17]. Alternatively, TE insertion can disrupt transcription factor binding sites (TFBS) within pre-existing CREs, thus attenuating or completely repressing nearby gene expression [25]. Additionally, insertion of TEs into coding sequences of genes may disrupt the open reading frames, and modify splicing sites [26–28]. Dysregulated gene expression due to TE insertions can lead to disease phenotypes as TE de-repression (i.e. de-methylation of TE sequences) is correlated with many neurological disorders and is a hallmark of multiple cancer types [29–40].

Over the last decade, the scientific community has begun to characterize the biological determinants of TE co-option in mammalian regulatory networks [8,13,14,17,20,41]. SINE-VNTR-Alus (SVAs) are the youngest human TEs. These transposons are composed of a 5’ CCCTCT hexamer repeat, an Alu-like element, a variable number of tandem repeats (VNTR), a SINE element derived from an ancestral endogenous retrovirus (HERVK-10), and a poly-A tail [42]. Six main SVA subfamilies have been characterized (SVA-A through -F) and nearly half of the annotated ∼3,700 copies are human-specific, including all the SVA-Es and -Fs.

Importantly, the SVAs are still actively transposing in the human genome by taking advantage of the L1-LINE machinery, and are among the most epigenetically de-repressed and transcriptionally upregulated TEs across a multitude of cancers and neurological disorders [17,24,26,33,43]. Given their young evolutionary age, SVAs provide a unique opportunity to elucidate how the human genome is evolving.

We, and others, have demonstrated that SVAs are frequently co-opted as functional enhancers and promoters in human and chimpanzee gene regulatory networks [16,17,20,24]. Yet we have an incomplete understanding of the extent of this evolutionary process and the underlying mechanisms. Why are some SVAs recruited as functional enhancers and others are not? What are the molecular properties underlying the regulatory potential of SVAs? Here, we answered these questions by exploring the highly permissive genomic environment of induced pluripotent stem cells (iPSCs) and NCCIT cells. The latter is a human cell line derived from an embryonal carcinoma which exhibits genomic properties comparable to human embryonic stem cells and iPSCs, including widespread de-methylation. These cells are particularly favorable for the study of transposable elements [18]. Compared to iPSCs, NCCITs are easier to manipulate and more suitable to perform Cas9-mediated genome editing experiments.

We demonstrate that ∼750 SVAs are depleted of repressive histone marks (i.e. H3K9me3) in iPSCs and in NCCITs. We found that the transcriptionally active SVAs are significantly closer to genes and harbor more TFBS than those harboring the repressive H3K9me3 epigenetic modification. Moreover, when comparing the sequence of the de-repressed SVAs with the repressed SVAs, we detected an enrichment for a long DNA motif composed of flanking binding sites for YY1/2 and OCT4 within the de-repressed group. The former is a ubiquitously expressed transcriptional regulator, while the latter is an essential regulator of cell pluripotency. Further the de-repressed SVAs are also enriched for individual (i.e. non-flanking) YY1 and OCT4 binding sites. We used Chromatin Immunoprecipitation followed by sequencing (ChIP-seq) to demonstrate that YY1 and OCT4 bind adjacently on many of the active SVAs. We leveraged CRISPR-interference (CRISPRi) to epigenetically repress active SVAs resulting in a loss of YY1 and OCT4 binding leading to massive alterations in gene expression.

## Results

### Human-specific SVAs are enriched in areas of de-repressed chromatin

We took advantage of publicly available H3K9me3 ChIP-seq data (paired-end long reads) generated in iPSCs [19] to assess to what extent SVAs are repressed in a pluripotent context. We only retained uniquely mapping high quality reads (Samtools q10 filtering). This analysis revealed that of all the SVAs annotated in the human genome (N=3,734; hg19 assembly), approximately 80% (2,983) are decorated with this repressive histone methylation in iPSCs (Fig. 1A). Conversely, 751 SVAs were depleted of H3K9me3, indicating that they are de-repressed (Fig. 1A).

**Figure 1.**
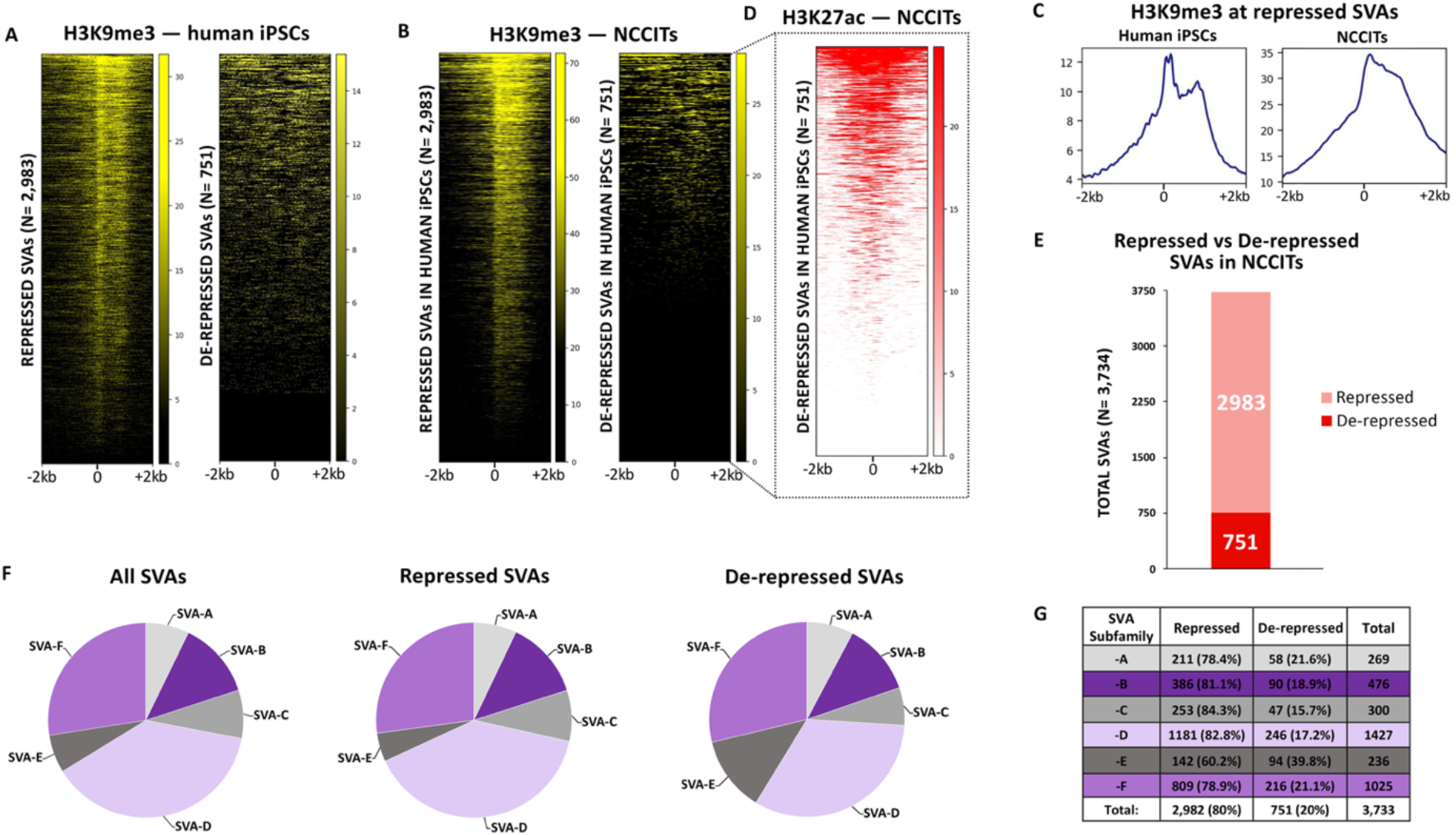
751 human SVAs are active in human pluripotent cells. (A) Heatmaps depicting H3K9me3 ChIP-seq signal at all human SVAs in iPSCs. (B) ChIP-seq for H3K9me3 in human NCCITs at SVA regions previously classified as repressed and de-repressed in human iPSCs. (C) Average profiles of H3K9me3 enrichment at SVAs repressed in human iPSCs and NCCITs. (D) ChIP-seq for H3K27ac in NCCIT cells. The heatmap is centered on the 751 de-repressed SVAs. (E) Barplot representing the number of total repressed and de-repressed human SVAs (i.e. SVAs decorated with H3K9me3 versus lacking H3K9me3). 2982 of the ∼3700 human SVAs were repressed, while 751 were de-repressed. (F) & (G) Human-specific SVAs, subfamilies SVA-E and -F are enriched within the de-repressed SVA population.

Next, we wanted to compare the iPSCs findings with data generated in NCCITs in our laboratory. As mentioned, NCCITs have pluripotent characteristics but compared to human iPSCs and embryonic stem cells, the NCCITs are significantly easier to manipulate and have high transfection efficiency, which makes them suitable for CRISPR experiments. We thus performed ChIP-seq for H3K9me3 to profile repressed SVA regions in NCCITs. To ensure high-quality read mappability of repetitive regions we sequenced 100 bp long paired-end reads and upon mapping, only retained the uniquely mapping high quality reads (Samtools q10 filtering). Notably, the SVA methylation pattern previously observed in iPSCs was perfectly recapitulated in NCCITs (Fig. 1B & 1C). These findings indicate that NCCITs share a similar TE epigenetic landscape with iPSCs making them a suitable model to study stem cell genomics, and support that 751 SVAs are de-repressed in pluripotent environments (hereafter “de-repressed SVAs”; Supplementary File S1).

We wanted to ensure that the lack of H3K9me3 signal on the de-repressed SVAs was not a consequence of mapping limitations, in that the youngest SVAs may be too similar to one another to allow for unique read mapping. To test this hypothesis, we looked at another histone modification. Specifically, we took advantage of publicly available H3K27ac ChIP-seq data (100bp paired-end) generated in NCCITs by the Wysocka laboratory [18]. H3K27ac usually decorates active cis-regulatory elements and we surmised that de-repressed SVAs may be transcriptionally active and therefore should exhibit ChIP-seq signal for this histone mark. This analysis revealed that the large majority of the de-repressed SVAs (636/751) are decorated with H3K27ac, which suggests they are in an active regulatory state, and that they are “mappable” with uniquely mapped reads (hereafter “active SVAs”; Fig. 1D). Additionally, 120 SVAs were not marked with H3K9me3 or H3K27ac. We attribute this pattern to potential mappability issues, and did not consider this SVA subset for downstream analyses. Overall, these data suggest that approximately 20% of all SVAs are located within transcriptionally de-repressed and active chromatin in iPSCs and in stem cell-like NCCITs (Fig. 1A-E).

SVAs can be further categorized into six evolutionarily conserved subfamilies (SVA-A through -F), two of which (SVA-E, -F, plus the F1 subgroup) are human-specific. We investigated the subfamily composition of the de-repressed and repressed SVAs and observed that human-specific SVA subfamilies -E and -F (including F1) were significantly enriched in the de-repressed group (Fisher’s Exact Test p < 1.0 ×10^−5^; Fig. 1F). In particular, ∼40% of the SVA-Es were found in a de-repressed state, as compared to an average of ∼19% for all the other families (Fisher’s Exact Test p < 2.2 × 10^−16^; Fig. 1G). In summary, these findings indicate that young, human-specific SVAs are enriched not only in de-repressed chromatin, but also in active regions in NCCIT cells.

### Sequence location and composition underlie SVA activation

We aimed to investigate the specific genomic features underlying the selective de-repression of SVAs. First, we reasoned that active SVAs preferentially located near coding genes could be co-opted as active cis-regulatory elements. We used the Genecode_v33 annotations and calculated the distance from the nearest transcription start site (TSS) for both repressed and active SVAs. We observed that active SVAs are significantly closer to coding genes when compared to the repressed SVAs (Wilcoxon’s Rank Sum Test p < 2.2 × 10^−16^; Fig. 2A). Namely, the active SVAs are located approximately 10 kb closer to the nearest TSS than the repressed SVAs (Fig. 2A).

**Figure 2.**
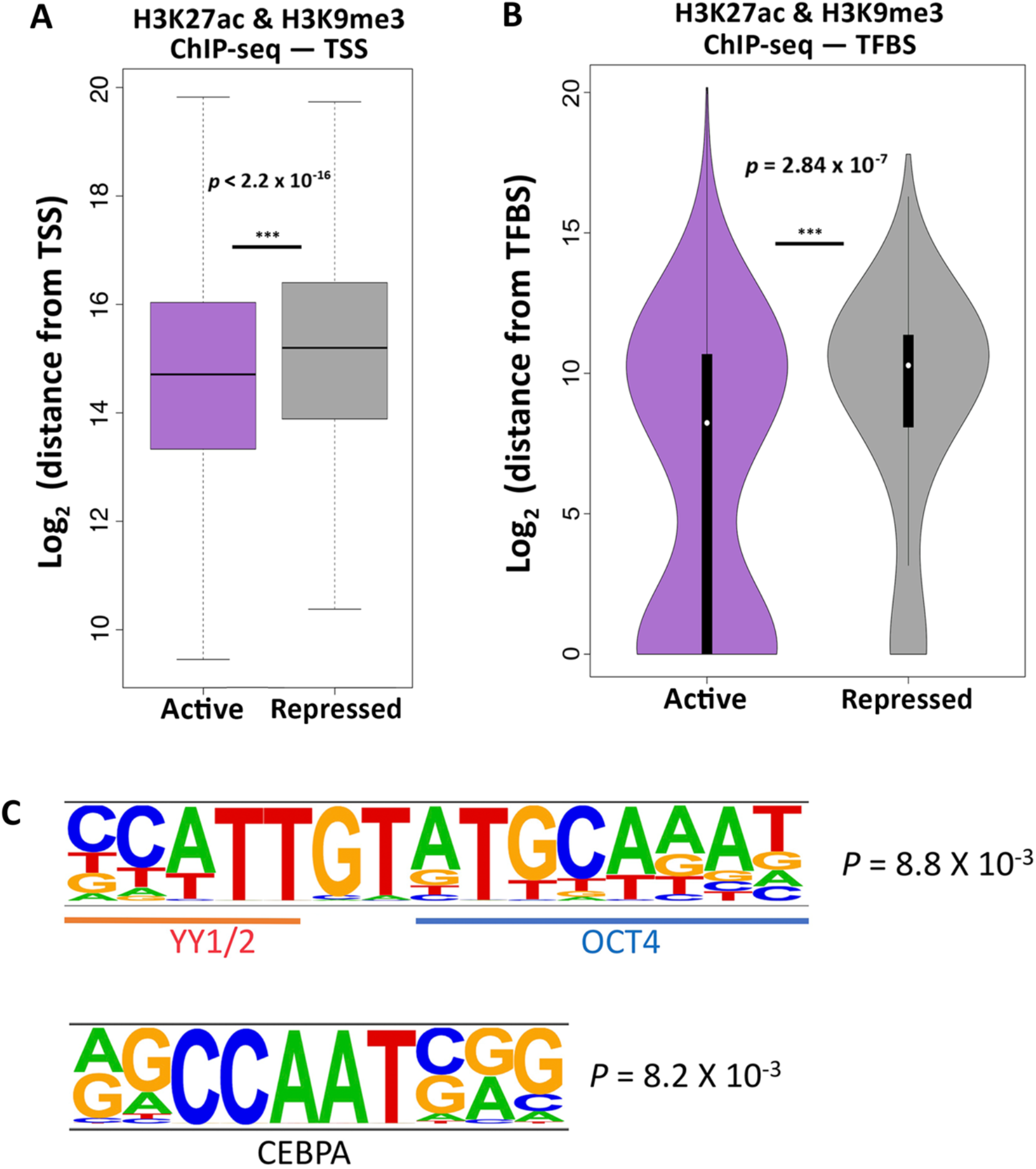
Specific genomic features characterize the de-repressed SVAs. (A) SVAs in an active configuration are approximately 10 kb closer to TSS than SVAs in a repressed configuration (Wilcoxon’s Rank Sum Test p < 2.2 × 10^−16^). (B) Active SVAs are significantly closer to, and directly overlap with, TFBS three times more than repressed SVAs (Wilcoxon’s rank-sum test p = 2.84 × 10^−7^). (C) Motif analysis performed on the de-repressed SVAs shows enrichment for consecutive YY1/2 and OCT4 motifs. A CEBPA motif was also enriched.

Next, we surmised that SVA de-repression may be a consequence of gene regulatory activity. Consistent with this hypothesis, we found that relative to the repressed SVAs, active SVAs were significantly closer to TFBS as defined by the Encode Consortium (Wilcoxon’s rank-sum test p = 2.84 × 10^−7^; Fig. 2B). Further, the active SVAs directly overlap a TFBS three times more frequently than the repressed SVAs (Fig. 2B). These data suggest that the active SVAs have a higher likelihood of exhibiting gene-regulatory activity.

To further explore this hypothesis, we performed sequence-based computational motif analysis on the de-repressed SVAs using HOMER with the repressed SVAs as the background control. With this approach, we identified an enrichment for a long motif composed of flanking YY1/2 and OCT4 binding sites in the de-repressed SVAs (Fig. 2C). YY1/2 are ubiquitously expressed transcriptional regulators, while OCT4 is one of the transcription factors essential for pluripotency maintenance. The YY1-OCT motif is found seven times more frequently in the de-repressed SVAs than in the repressed ones. Importantly, not only is the flanking YY1-OCT4 motif enriched, but also individual (i.e. non-flanking) YY1 and OCT4 motifs are enriched in the de-repressed SVAs as compared to the repressed ones. For instance, OCT4 alone is found in 12% of the de-repressed SVAs, as opposed to 1% of repressed SVAs.

Additionally, we identified an enriched CEBPA motif (Fig. 2C). We conducted the opposite analysis, by looking for motifs enriched in repressed SVAs (using de-repressed as a background) and found several enriched motifs, including: SMAD3, KLF5 and several Krüppel-associated box (KRAB) domain-containing zinc-finger proteins (KZFPs).

### Repression of conventionally de-repressed SVAs has global consequences on gene expression

Since our results indicate that ∼20% of all SVAs may act as functional CREs in NCCITs, we aimed to validate our ChIP-seq data and computational predictions with a functional approach. We utilized CRISPRi to repress the 751 de-repressed SVAs. We leveraged the piggyBac transposon system (Systems Bioscience) to generate NCCIT cells with tetracycline-inducible expression of a functionally dead Cas9 (dCas9) fused to a repressive KRAB domain. We subsequently knocked-in two previously validated [20] single guide RNAs (sgRNAs) that simultaneously target approximately 80% of all the SVAs annotated in the human genome (Fig. 3A). The sgRNAs target the dCas9-KRAB to de-repressed SVAs leading to the deposition of the repressive histone methylation H3K9me3 by the KRAB domain (Fig. 3A). This NCCIT cell line exhibiting tetracycline-inducible dCas9-KRAB expression and constitutive dual sgRNA expression is hereafter referred to as SCi-NCCITs (i.e. SVA-CRISPRi-NCCITs). We treated the SCi-NCCITs with doxycycline for 72 hours to robustly induce dCas9-KRAB activation and function (Fig. 3B). Activation of dCas9-KRAB led to the accumulation of repressive H3K9me3 modifications on 620 of the 751 SVAs originally de-repressed in NCCITs (Fig. 3C & 3D). Importantly, this further demonstrates that reads can be uniquely mapped on this group of SVAs.

**Figure 3.**
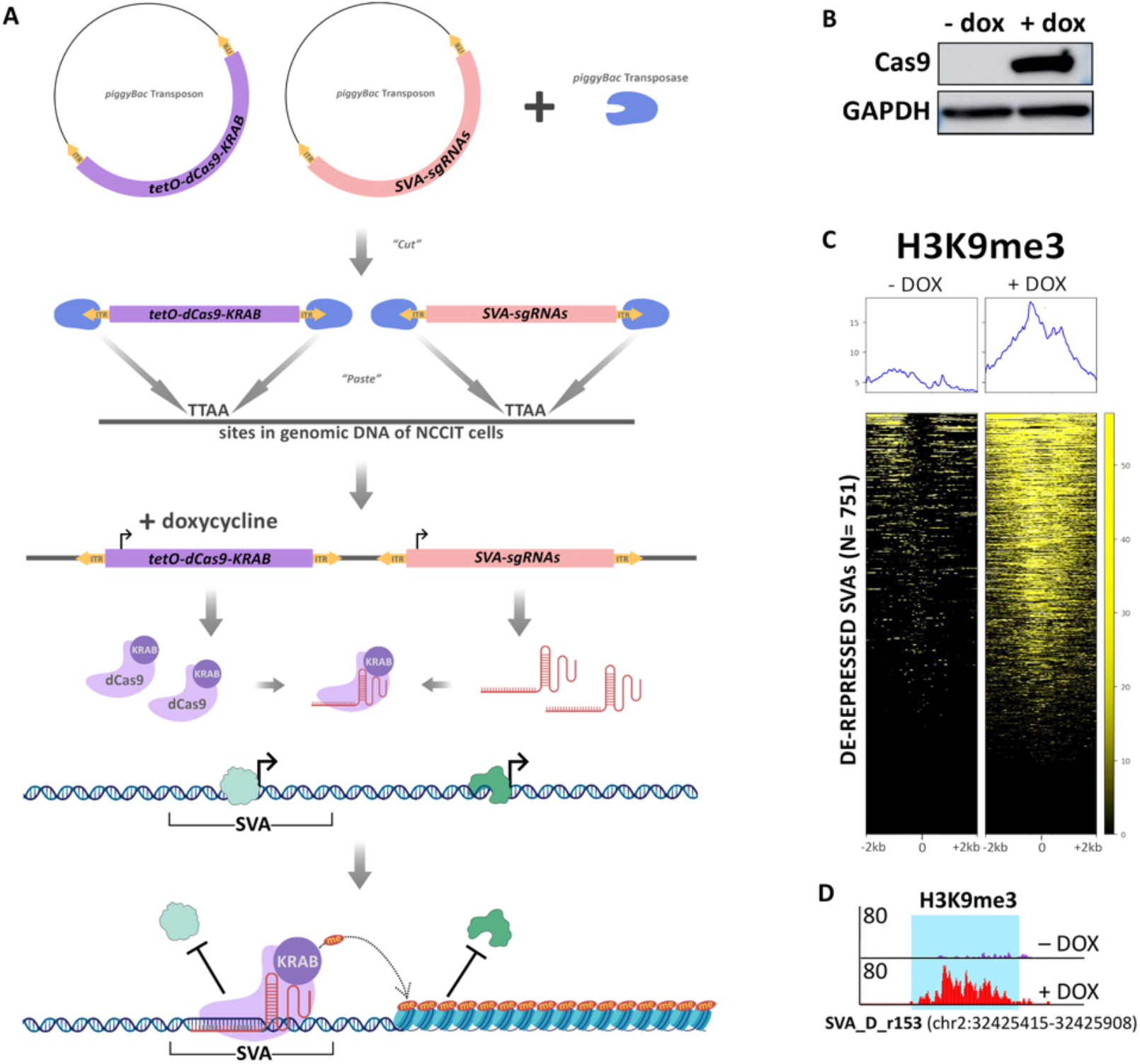
Epigenetic manipulation of NCCITs via CRISPRi results in SVA repression. (A) Schematic of the generation of the SCi-NCCIT line (created with BioRender.com). (B) Cas9 immunoblot displaying activation of dCas9-KRAB 72 hours post-doxycycline induction. (C) H3K9me3 ChIP-seq heatmap of the SCi-NCCITs shows increased H3K9me3 signal post-doxycycline treatment. (D) Genome browser screenshot displaying increased H3K9me3 before and after treatment with doxycycline at a de-repressed SVA-D.

To assess the impact of global SVA repression on genome-wide gene expression, we performed RNA-sequencing on the SCi-NCCITs in the presence and absence of doxycycline treatment (three replicates per condition). We identified 3,085 genes as differentially expressed upon induction of SVA repression (FDR < 0.05; Supplementary File S2) of which 1,596 were identified as upregulated and 1,489 as downregulated (Fig. 4A). The sgRNAs used for this experiment were originally designed to target a DNA sequence shared by SVAs with the LTR5Hs family [20]. Given this premise, we restricted our analysis exclusively to genes putatively associated with human SVAs (i.e. considering only the genes that represent the closest gene to an annotated SVA). Overall, 131 of the 3,085 differentially expressed genes represented the closest gene to an SVA (Fig. 4B), 101 of which were specifically near a de-repressed SVA. This number is significantly higher than expected by chance (Fisher’s Exact Test p < 0.001), suggesting that the expression of most of these 131 genes is likely under the direct control of SVAs. Importantly, 109 of the 131 SVA-regulated genes (83.2%) were downregulated upon doxycycline treatment (Fig. 4B), indicating that SVA de-repression is necessary for gene activation.

**Figure 4.**
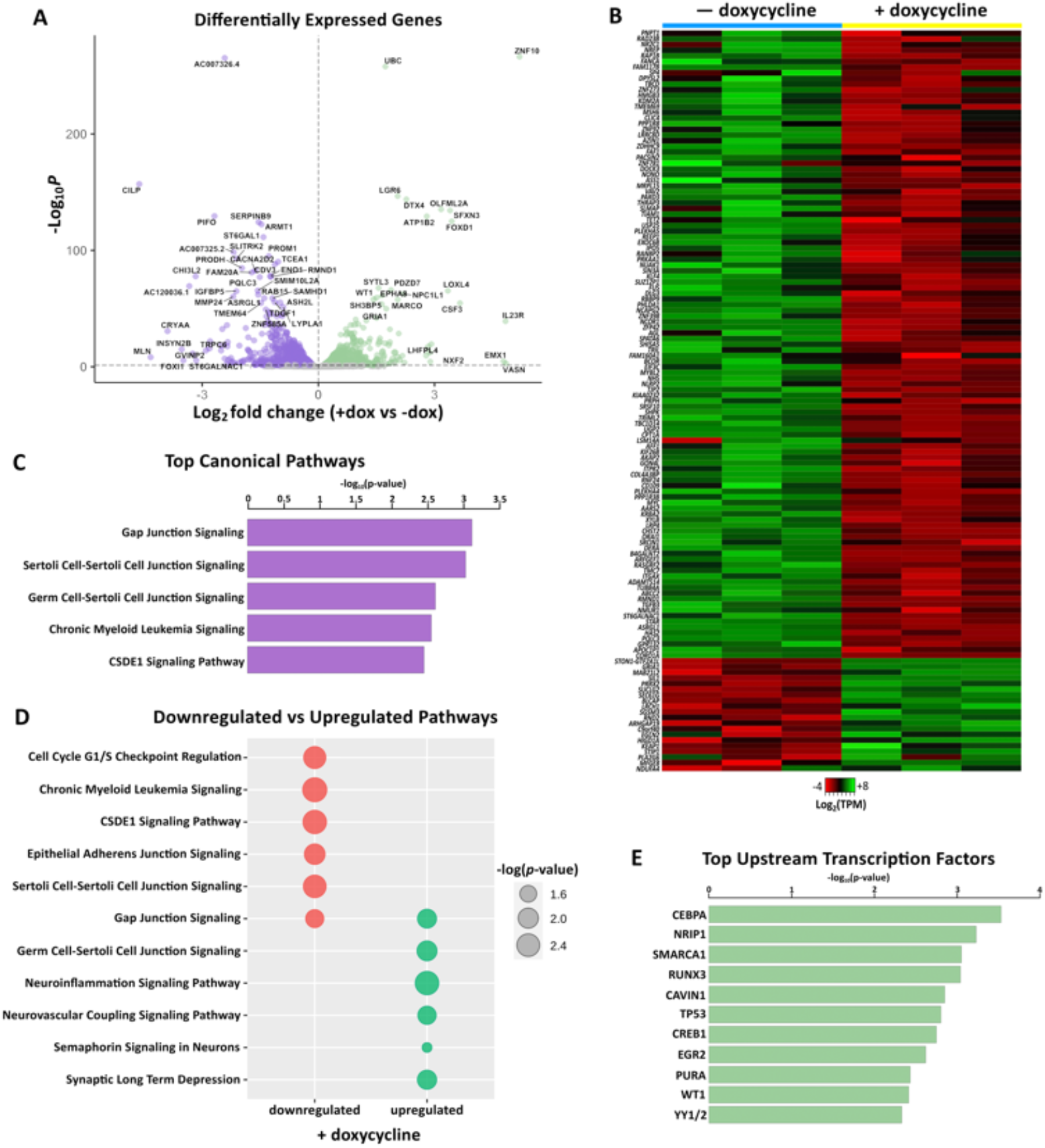
CRISPRi-mediated repression of conventionally de-repressed SVAs results in aberrant gene expression. (A) Volcano plot showing genes differentially expressed in SCi-NCCITs after doxycycline treatment (purple = downregulated genes; green = upregulated genes). (B) Heatmap of 131 genes that are differentially expressed post-doxycycline treatment and also represent the nearest gene to an SVA. (C) Top canonical pathways predicted by IPA for the 131 genes differentially expressed after doxycycline treatment. (D) Canonical pathways predicted by IPA for the 111 downregulated (red) and 20 upregulated (green) genes. (E) Top upstream regulators/transcription factors predicted by IPA for the 131 genes differentially expressed after doxycycline treatment.

We then assessed if any of the differential gene expression is due to repressing LTR5Hs since the sgRNAs also target DNA sequences in this TE family. Only 120/3,085 differentially expressed genes represented the closest gene to an LTR5H. Additionally, we detected very minimal overlap (8 genes in total) between the differentially expressed genes near LTR5Hs and the differentially expressed genes near SVAs. These analyses ensure that the alterations in gene expression are due to the repression of SVAs and not LTR5Hs. In summary, these data indicate that the SVAs directly regulate the expression of at least 130 genes and that SVA repression may contribute to the differential expression of up to ∼2,800 genes, potentially as a cascade effect.

We performed Ingenuity Pathway Analysis (IPA) on the 131 SVA-regulated genes and found an enrichment for gap junction signaling processes including Gap Junction Signaling, Sertoli Cell-Sertoli Cell Junction Signaling, and Germ Cell-Sertoli Cell Junction Signaling (Figs. 4C, D). These processes are important during gametogenesis and given that NCCIT cells retain pluripotency-like characteristics, we speculate that co-opted SVAs could play a role in regulating germ line development and gametogenesis. However, it is important to note that the NCCIT cell line is derived from an embryonal testicular carcinoma, and this could bias our IPA. Nonetheless, recent studies have corroborated an SVA contribution in germ cells and showed that in human primordial germ cells SVA sites are largely hypomethylated and genes proximal to SVAs are expressed at a higher level compared to embryonic stem cells [20,44,45]. Additionally, SVA transcription and retrotransposition are specifically seen in both human spermatozoa [46] and oocytes [47].

We next used IPA to look for top upstream transcriptional regulators of the 131 SVA-associated genes. This analysis identified transcription factors previously highlighted by the motif analyses (CEBPA and YY1/2), along with several others including SMARCA1, CREB1, and WT1 (Fig. 4E). While YY1 is an essential ubiquitous developmental regulator [48], CEBPA is a regulator of differentiation in the hematopoietic [49] and adipocyte [50] lineages, and is involved in gametogenesis. SMARCA1 is a chromatin remodeler [51], CREB1 plays a role in both steroidal [52] and non-steroidal [53] female hormonal stimulation, and WT1 is involved in urogenital specification distinctively associated with the differentiation of Sertoli cells [54]. These data suggest a possible SVA contribution to germ line developmental processes and demonstrates the genome-wide impact of SVA repression on gene expression.

### YY1 and OCT4 mediate SVA regulatory activity

Since our motif analysis revealed that de-repressed SVAs are enriched for adjacent YY1/2-OCT4 binding motifs and given that many differentially expressed genes were YY1/2 targets, we performed ChIP-seq for YY1 and OCT4 in SCi-NCCITs with and without doxycycline treatment. As done previously for the histone modification ChIP-seq, we sequenced 100 bp long paired-end reads to ensure high mappability efficiency.

These experiments revealed that under normal conditions (i.e. no doxycycline treatment) YY1 and OCT4 bind 288 and 54 SVAs, respectively (Figs. 5A & 5B). Specifically, we identified two different clusters of YY1 binding: one located in the *Alu*-like region (Fig. 5A top cluster), and a second near the start of the SINE element, likely in proximity of the HERVK10-derived promoter in the SINE region (Fig. 5A second cluster). OCT4 binding was limited to this second region, whereas we did not detect binding for this transcription factor in the *Alu*-like region (Fig. 5B). The SVAs bound by YY1 and OCT4 were all de-repressed SVAs decorated with H3K27ac. Importantly, binding of both YY1 and OCT4 was significantly attenuated upon SVA repression via CRISPRi (Figs. 5A, B). We observed a significant overlap with 33 SVAs bound by both YY1 and OCT4 in the SINE region. In this case, the binding of the two transcription factors was sequential (i.e. one next to each other) as originally predicted by our motif analysis (Fig. 5C). This pattern was found in de-repressed SVAs of all the main families (SVA-A through -F).

**Figure 5.**
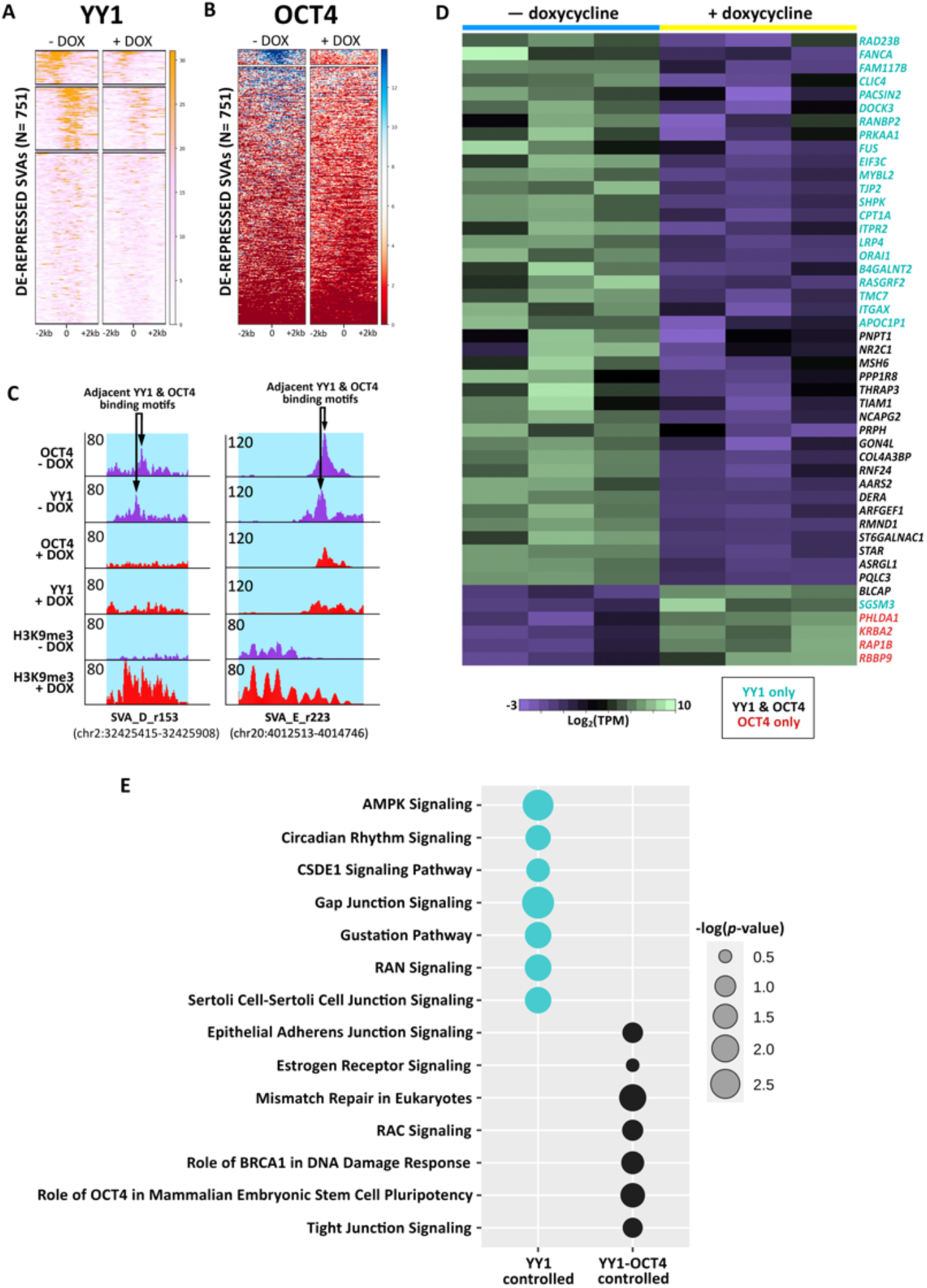
Individual and sequential binding of YY1 and OCT4 contributes to SVA regulatory activity. (A) YY1 ChIP-seq signal at the 751 de-repressed SVAs before and after doxycycline treatment. (B) OCT4 ChIP-seq signal at the 751 de-repressed SVAs before and after doxycycline treatment. (C) Genome browser screenshot displaying the sequential binding of YY1 and OCT4 at a truncated, de-repressed SVA-D and a full-length, de-repressed, human-specific SVA-E. Upon doxycycline induction, the binding of both transcription factors is lost, while H3K9me3 signal is gained. The full length SVA-E is located near the gene *RNF24* (specifically 18kb from the TSS), which is downregulated upon CRISPRi mediated SVA repression (logFC =-0.56; Adjusted p=9.14 × 10^−11^) (D) Heatmap of 47 genes that are differentially expressed upon doxycycline treatment and regulated by YY1 and/or OCT4 (Black: YY1 and OCT4, Teal: YY1 only, Red: OCT4 only). (E) Canonical pathways predicted by IPA for the SVA-regulated genes that are differentially expressed upon doxycycline treatment and regulated by YY1-only (teal) and YY1-OCT4 (black).

Finally, we leveraged the nearest TSS approach to determine the closest gene to each of the SVAs bound by YY1 and/or OCT4. Using our RNA-seq data, we investigated whether loss of YY1 and OCT4 binding altered the expression of neighboring genes. Notably, 44 of 288 genes located near YY1-bound SVAs and 24 of 54 genes located near OCT4-bound SVAs were differentially expressed upon CRISPRi-induced SVA repression (Fig. 5D). Interestingly, of the 6 genes upregulated upon doxycycline treatment, 4 were located near SVAs bound exclusively by OCT4 (Fig. 5D). Overall, 85% of the genes (28/33) located near SVAs that were bound by both YY1 and OCT4 were differentially expressed upon CRISPRi-induced SVA repression (Fig. 5D). This suggests that synergistic binding of these two transcription factors on the SVA sequence is important for the regulatory activity driven by these transposable elements (Fig. 5D & Fig. 6). Pathway Analysis revealed that the genes regulated by the cooperative YY1-OCT4 binding (and by OCT4-only) are enriched for processes related to pluripotency and DNA repair (Fig. 5E). Conversely, the genes bound by YY1-only are enriched for processes related to germline development (Fig. 5E).

**Figure 6.**
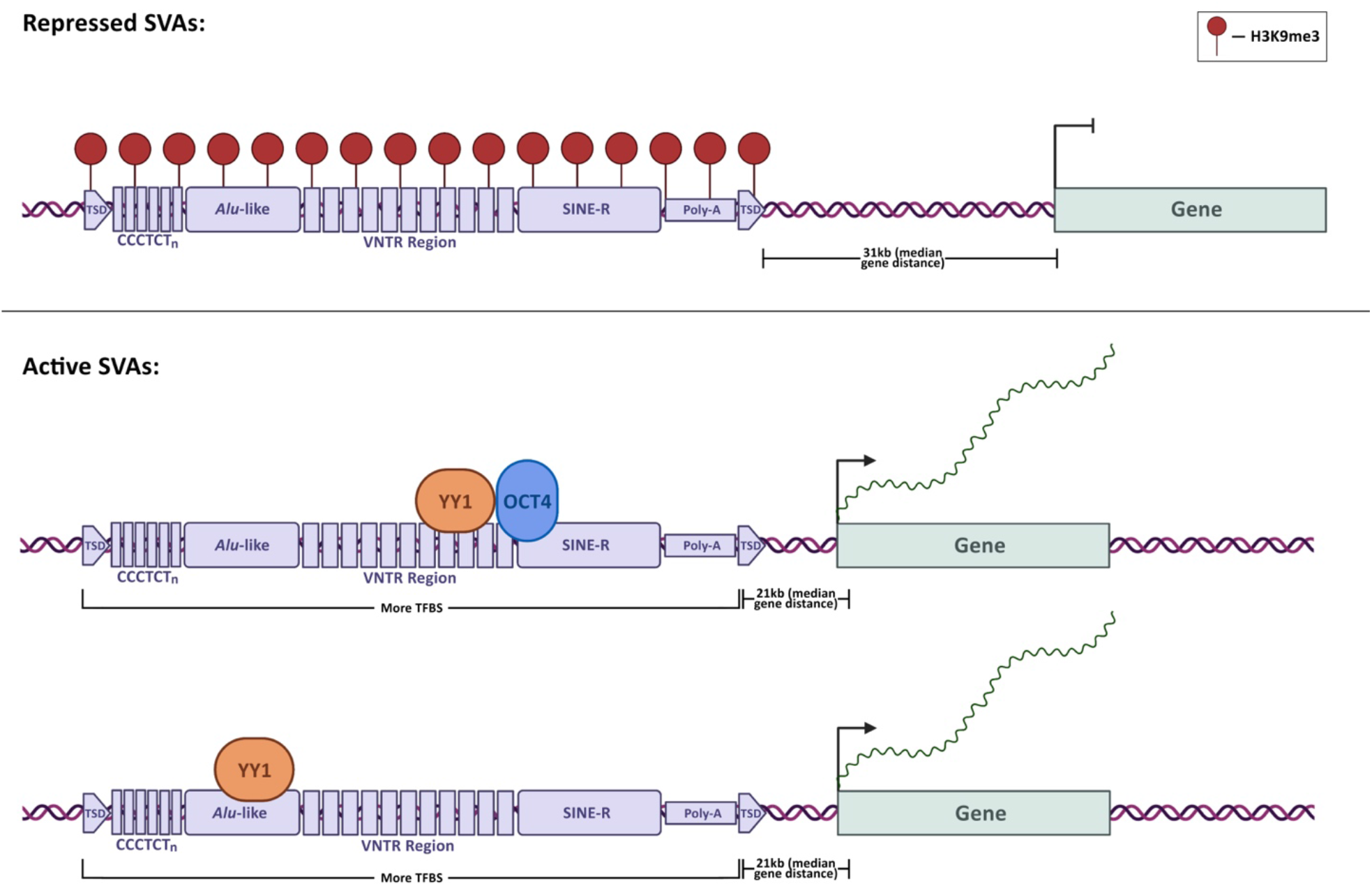
Model for SVA co-option in human pluripotent stem cell gene regulation. This figure was created with BioRender.com.

## Discussion

SVAs are evolutionarily young transposable elements, which colonized the great ape genomes in the last 10-15 million years. In fact, they are not found in gibbons whose lineage split from the remaining apes ∼17 million years ago [55]. Interestingly, the gibbon genome has been independently colonized by a distinct family of transposons called LAVAs (LINE-AluSz-VNTR-Alu) [56,57]. Notably, SVA and LAVA structures are similar, and both have shown high cis-regulatory potential and massive co-option into gene regulatory networks [16,17,20,57]. However, the mechanisms underlying SVA and LAVA domestication and recruitment into primate gene regulatory networks have not been explored in depth.

Here, we aimed at elucidating such mechanisms, focusing specifically on SVAs and stem cells. We show that ∼20% of all the human SVAs are found in a de-repressed chromatin state in iPSCs and in NCCITs, with a near perfect overlap between the two cell types. Most of these de-repressed SVAs were also decorated with histone modifications characteristic of active enhancers and promoters (H3K27ac), suggesting that their de-repression is associated with cis-regulatory activity. This pattern may indicate that either the SVAs are enriched in regions that were already transcriptionally active before their insertion, or that the SVAs themselves dictated the epigenetic landscape as a consequence of their sequence. A body of literature has emerged supporting the contribution of specific TE families to the dispersion of TFBS and cis-regulatory elements in the eukaryotic genomes (Fueyo et al., 2022). This can happen in multiple ways: TEs already harbor the TFBS before the transposition event, or they gain TFBS afterwards as a consequence of new mutations. In the former case, if TEs harboring TFBSs insert near genes, this will increase the likelihood for the TE to be co-opted as an enhancer/promoter and thus lose repressive H3K9me3 and gain H3K27ac. Consistent with this scenario, it is estimated that in human embryonic stem cells ∼20% of the TFBSs for pluripotency factors are located within transposable elements (Kunarso et al. 2010; Sundaram et al 2017; Fueyo et al. 2022).

The youngest SVAs (which are human-exclusive), and especially the SVA-Es, are the most enriched among the active copies. To this end, we speculate that the older SVAs accumulated genetic mutations over time, hampering their regulatory potential. Alternatively, the human genome may be adapted to silence older transposons, and this may have affected copies with higher regulatory potential.

According to our data, SVA location and sequence composition are the best predictors of cis-regulatory activity. Active SVAs are, on average, 10 kb closer to genes than the repressed ones. This is likely due to the fact that the chromatin environment near gene loci may be more frequently in an accessible and de-repressed state and is thus more suitable for co-option of novel cis-regulatory elements. Moreover, a shorter distance from the nearest TSS may facilitate the tri-dimensional interaction between the SVA-derived enhancer and the gene promoter. This is also in line with a recent study that demonstrated that eQTL-rich TEs tend to be significantly closer to genes than eQTL-poor TEs [25].

We show that active SVAs host a significantly higher number of TFBS than the repressed ones. This coincides with many studies suggesting that TE exaptation into gene regulatory networks is largely driven by the evidence that they propagate TFBS across the genome [60]. We demonstrate that a long motif with flanking YY1/2-OCT4 binding sites is enriched in de-repressed SVA copies relative to repressed SVAs. This may mediate their function in gene regulatory networks of stem and stem-like cells. In fact, OCT4 is one of the four Yamanaka factors [61] essential for pluripotency maintenance. YY1 is one of the major transcriptional regulators in human cells; it is ubiquitously expressed across all cell-types and performs many different functions in transcriptional regulation, including transcriptional activation, repression, as well as mediation of enhancer-promoter looping [62]. To this regard, over 40% of the SVAs de-repressed in iPSCs and NCCITs are bound by YY1, OCT4, or both. Importantly, when both are present, they bind alongside each other as predicted by computational motif analysis. As expected, depositing repressive histone methylation (H3K9me3) on the SVA locus nearly abolishes the binding of these two transcription factors near SVAs, leading to an alteration of nearby gene expression. In fact, even when restricting the analysis to the genes that are located near SVAs, repressing 620 of the 751 SVAs normally active in NCCITs results in 131 of differentially expressed genes. The expression of most (88%) of these genes is attenuated with the repression of the nearby SVA, further supporting the enhancer activity provided by these transposons. These genes include important regulators of cell pluripotency and cell differentiation, such as *MYC, MYBL2, FUS, ITGAX, SP4* and several others. We remark that this analysis was very conservative. In fact, given the nature of the sgRNAs that we chose, which target both SVAs and LTR5Hs, we exclusively focused on the nearest gene to an SVA transposon, and thus the number of genes affected by distal SVA repression is likely much higher.

Finally, repressing SVAs did not result in any obvious alteration in cellular phenotypes, although we cannot rule out that a longer experiment (i.e. with cells collected more than 72 hours post doxycycline treatment) may result in alterations in cell viability/proliferation as a consequence of SVA-repression. Future studies may assess longer term SVA repression, focusing on the ability of iPSCs to act as pluripotent cells upon sustained repression of the SVA-derived enhancers. In summary, in this study we provide further evidence that SVA transposons are an important component of the human gene regulatory networks, specifically in stem and stem-like cells. We propose a potential mechanism underlying this cis-regulatory activity where SVA location and sequence composition regulate this co-option. Additional studies are required to determine if the YY1-OCT4 combination is a driver of SVA regulatory activity only in pluripotent cells or, alternatively, in a broader, more universal context. In this study most genomics experiments were conducted in NCCIT cells. Further genomic studies in human iPSCs will be necessary to confirm the proposed mechanism.

## Materials and Methods

### Antibodies and sgRNAs

YY1 ChIP-seq: Cell Signaling Technology D5D9Z/46395S (15ug per ChIP). OCT4 ChIP-seq: Abcam ab181557 (15ug per ChIP). H3K27me3 ChIP-seq: Abcam ab8898 (3ug per ChIP). Cas9 Western Blot: Active Motif 61757 (1:100). GAPDH Western Blot: Cell Signaling Technology D16H11/5174 (1:1000). Anti-Rabbit IgG, HRP-linked Western Blot: Cell Signaling Technology 7074 (1:10000). Anti-Mouse IgG, HRP-linked Western Blot: Cell Signaling Technology 7076 (1:10000). The two sgRNAs were designed and used in a previous study [20]: sgRNA1: 5’ CTCCCTAATCTCAAGTACCC 3’ ; sgRNA2: 5’ TGTTTCAGAGAGCACGGGGT 3’.

### NCCIT Cell Line Culture

The NCCIT cell line (ATCC) was maintained in RPMI media supplemented with 10% tet-free FBS, 1% penicillin-streptomycin solution, and 1% L-glutamine and incubated at 5% CO2, 20% O2 at 37°C.

### ChIP-Sequencing

All samples from different conditions were processed together to prevent batch effects. Between 10-15 million cells were cross-linked with 1% formaldehyde for 5 minutes at room temperature, quenched with 125 mM glycine, harvested, and washed twice with 1x PBS. The fixed cell pellet was resuspended in ChIP lysis buffer (150 mM NaCl, 1% Triton X-100, 0.7% SDS, 500 μM DTT, 10 mM Tris-HCl, 5 mM EDTA) and chromatin was sheared to an average length of 200–900 base-pairs, using a Covaris S220 Ultrasonicator. The chromatin lysate was diluted with SDS-free ChIP lysis buffer. 15μg of antibody was used for YY1 and OCT4 and 3μg of antibody for H3K9me3. The antibody was added to 5μg of sonicated chromatin along with Dynabeads Protein G magnetic beads (Invitrogen) and incubated at 4°C overnight. The beads were washed twice with each of the following buffers: Mixed Micelle Buffer (150 mM NaCl, 1% Triton X-100, 0.2% SDS, 20 mM Tris-HCl, 5 mM EDTA, 65% sucrose), Buffer 200 (200 mM NaCl, 1% Triton X-100, 0.1% sodium deoxycholate, 25 mM HEPES, 10 mM Tris-HCl, 1 mM EDTA), LiCl detergent wash (250 mM LiCl, 0.5% sodium deoxycholate, 0.5% NP-40, 10 mM Tris-HCl, 1 mM EDTA) and a final wash was performed with 0.1X TE. Finally, beads were resuspended in 1X TE containing 1% SDS and incubated at 65°C for 10 min to elute immunocomplexes. The elution was repeated twice and the samples were incubated overnight at 65°C to reverse cross-linking, along with the untreated input (5% of the starting material). The DNA was digested with 0.5 mg/ml Proteinase K for 1 hour at 65°C and then purified using the ChIP DNA Clean & Concentrator kit (Zymo) and quantified with QUBIT. Barcoded libraries were made with NEBNext Ultra II DNA Library Prep Kit for Illumina (New England BioLabs) and sequenced on an Illumina NextSeq 2000 producing 100 bp paired-end reads.

### ChIP-seq Analysis

After removing the adapters with TrimGalore!, the sequences were aligned to the reference hg19, using Burrows-Wheeler Alignment tool, with the MEM algorithm [63]. Uniquely mapping aligned reads were filtered based on mapping quality (MAPQ > 10) to restrict our analysis to higher quality and likely uniquely mapped reads, and PCR duplicates were removed. Peaks were called for each SVA site using the default parameters, at 5% FDR, with default parameters

### Generation and Culturing of SCi-NCCIT Stable Cell Lines

A plasmid with a tetracycline-inducible dCas9-KRAB expression cassette flanked by piggyBac recombination sites was obtained from the Wysocka Lab at Stanford University. This plasmid ‘p-dCas9-KRAB’ confers constitutive puromycin resistance, allowing for selection of stably transduced clones when co-expressed with the piggyBac transposase plasmid (‘p-PB-Transposase’, Systems Bioscience). The p-dCas9-KRAB and p-PB-Transposase plasmids were co-transfected into NCCIT cells (ATCC) at 70% confluency using a 6:1 ratio of Fugene HD (Promega) for 48 hours. Two days post-transfection, cells were treated with puromycin selective media at a concentration of 1 ug/mL. Stable clones were isolated and dCas9 expression assessed via Western blot. Next, we obtained a piggyBac transposon plasmid containing the two sgRNAs [20] targeting ∼80% of all annotated SVAs in humans termed ‘p-sgRNA’ (Systems Bioscience). This plasmid constitutively confers dual sgRNA expression and geneticin resistance. The p-sgRNA and p-PB-Transposase plasmids were co-transfected into the NCCIT-dCas9KRAB cells as above. Two days post-transfection cell were treated with geneticin selective media at a concentration of 400 ug/mL. Following antibiotic selection, the NCCIT-dCas9KRAB-SVAsgRNA (SCi-NCCITs) cell line was maintained in ATCC-formulated RPMI media supplemented with 10% tet-free FBS, 1% L-glutamine, 1µg/mL puromycin, and 400 µg/mL geneticin and incubated at 5% CO2, 20% O2 at 37°C. The SCi-NCCITs were seeded to 40% confluency and treated with 2 ug/mL doxycycline (Sigma Aldrich) for 72 hours. For all molecular and genomic CRISPRi experiments, dCas9-KRAB expression was induced with doxycycline for 72 hours.

### Western Blot

Cells were washed three times in PBS and lysed in radioimmunoprecipitation assay buffer (RIPA buffer) (50 mM Tris-HCl pH7.5, 150 mM NaCl, 1% Igepal, 0.5% sodium deoxycholate, 0.1% SDS 500 μM DTT) with protease inhibitors. Approximately 40 μg of whole cell lysate were loaded in Novex WedgeWell 4–20% Tris-Glycine Gel (Invitrogen) and subject to SDS-PAGE. Proteins were then transferred to a Immun-Blot PVDF membrane (ThermoFisher) for antibody probing. Membranes were blocked with a 10% BSA in TBST solution for 30 minutes then incubated with primary antibodies in a 5% BSA in TBST, diluted as above. Next, membranes were washed with TBST and incubated with secondary antibodies, diluted as above. Chemiluminescent signal was detected using the Pierce ECL Plus Western Blotting Substrate (ThermoFisher) and an Amersham Imager 680.

### RNA Extraction and Library Preparation for RNA sequencing

Cells were lysed in Tri-reagent (Zymo) and total RNA was extracted using Direct-zol RNA Miniprep kit (Zymo) according to the manufacturer’s instructions. RNA was quantified using DeNovix DS-11 Spectrophotometer while the RNA integrity was checked on a Bioanalyzer 2100 (Agilent). Only samples with RIN value above 8.0 were used for transcriptome analysis. RNA libraries were prepared using 1μg of total RNA input using NEB-Next® Poly(A) mRNA Magnetic Isolation Module, NEBNext® UltraTM II Directional RNA Library Prep Kit for Illumina® and NEBNext® UltraTM II DNA Library Prep Kit for Illumina® according to the manufacturer’s instructions (New England Biolabs). Paired-end 100 bp reads were generated.

### RNA-seq Analysis

Reads were aligned to hg19 using STAR v2.567 [64], in 2-pass mode. Bam files were filtered based on alignment quality (q = 10) using Samtools [63]. We used the latest annotations obtained from Ensembl to build reference indexes for the STAR alignment. Adapters were removed with TrimGalore! and Kallisto [65] was used to count reads mapping to each gene. We analyzed differential gene expression with DESeq2 [66].

### Statistical and Genomic Analyses

All statistical analyses were performed using BEDTools v2.27.1 [67], DeepTools, and R v4.1.2. Fasta files of the regions of interest were produced using BEDTools v2.27.1 [67]. Shuffled input sequences were used as background. E-values < 0.001 were used as a threshold for significance. Motif analysis of de-repressed SVAs on a repressed SVA background was performed using HOMER [68]. Pathway analysis was performed with Ingenuity-Pathway Analysis Suite (Qiagen Inc., https://www.qiagenbioinformatics.com/products/ingenuity-pathway-analysis).

## Acknowledgements

The authors are grateful to Dr. Joanna Wysocka’s lab for providing the dCas9-KRAB piggyBac plasmid and in particular to Dr. Raquel Fueyo for providing critical support for the SCi-NCCIT CRISPRi line generation. The authors thank the Genomic Facility at The Wistar Institute (Philadelphia, PA) for the Next Generation Illumina Sequencing. We thank Dr. Geoffrey Faulkner (University of Queensland) and another anonymous reviewer for their insightful comments on our paper.

## Data availability

The original genome-wide data generated in this study have been deposited in the GEO database under accession code GSE192951.

## Author contributions

MTrizzino designed the project. SMB and AI generated the stable SCi-NCCIT CRISPR line; AI performed a preliminary set of genomic experiments. SMB performed all the genomic experiments. MTrizzino, SMB, and DTM analyzed the data. MTrizzino and SMB wrote the manuscript. CB provided essential advice and contribution for the planning of the CRISPR experiment. LP, SP and MTracewell contributed to some of the experiments. All the authors read and approved the manuscript.

